# Prefrontal cortex melanocortin 4 receptors (MC4R) mediate food intake behavior in mice

**DOI:** 10.1101/2022.06.01.494383

**Authors:** Rachel A Ross, Angela Kim, Priyanka Das, Yan Li, Yong Kee Choi, Andy T Thompson, Ella Douglas, Siva Subramanian, Kat Ramos, Kathryn Callahan, Vadim Y Bolshakov, Kerry J Ressler

**Author notes:** co-first author (Rachel A Ross, Angela Kim). corresponding author Rachel A. Ross, MD, PhD, Albert Einstein College of Medicine, 1410 Pelham Parkway S, Kennedy Building 118, Bronx, NY 10461.

## Abstract

**Background:** Melanocortin 4 receptor (MC4R) activity in the hypothalamus is crucial for regulation of metabolism and food intake. The peptide ligands for the MC4R are associated with feeding, energy expenditure, and also with complex behaviors that orchestrate energy intake and expenditure, but the downstream neuroanatomical and neurochemical targets associated with these behaviors are elusive. In addition to strong expression in the hypothalamus, the MC4R is highly expressed in the medial prefrontal cortex, a region involved in executive function and decision-making.

**Methods:** Using viral techniques in genetically modified mice combined with molecular techniques, we identify and describe the neuronal dynamics, and define the effects on feeding behavior of a novel population of MC4R expressing neurons in the infralimbic region of the cortex.

**Results:** Here, we describe a novel population of MC4R-expressing neurons in the infralimbic (IL) region of the mouse prefrontal cortex that are glutamatergic, receive input from melanocortinergic neurons of the arcuate hypothalamus, and project to multiple regions that coordinate appetitive responses to food-related stimuli. The neurons are depolarized by application of MC4R-specific peptidergic agonist, THIQ. Deletion of MC4R from the IL neurons causes increased food intake and body weight gain and impaired executive function in simple food-related behavior tasks.

**Conclusion:** Together, these data suggest that MC4R neurons of the IL play a critical role in the regulation of food intake.

## 1. Introduction

Food-seeking behavior is a complex process critical to survival, and it requires integration of multiple stimuli from different environmental and internal sensory modalities. Human studies using fMRI in response to food related stimuli have demonstrated that activity changes in medial aspects of the prefrontal cortex are correlated with body habitus[1], suggesting that food-related signals may influence cognitive function.

The medial prefrontal cortex (mPFC) regions in rodents, including prelimbic and infralimbic cortex, are critical for coordinating goal-directed and habit-based behavior[2]. The neurons in this region integrate sensory and limbic information to flexibly guide animal behavior based on learned information and in response to changes in inputs from other brain regions. These processes are essential to food-seeking and food-intake behavior, including meal termination. Accordingly, silencing of the mPFC in rats leads to hyperphagia of palatable food[3], likely due to reduced inhibitory control. Neurons within mPFC region have also been shown to be involved in driving food intake behavior in rodents[4], and responding to hunger state[5], but no studies have investigated the circuitry that allows this coordination of metabolic, cognitive, and motivated functions.

The melanocortin system is a well-known peptidergic network that processes food-related and hunger-state based signals within the brain. Mutations in the melanocortin 4 receptor (MC4R), one of the main melanocortinergic receptors expressed in the brain, are strongly associated with hyperphagia and obesity[6, 7]. In the central nervous system, MC4R is expressed in neuronal populations associated with regulation of metabolism, food intake, and food seeking behaviors[8]. Re-expression of MC4R in the paraventricular nucleus (PVH) in an MC4R knockout mouse, and deletion of MC4R from the PVH alone, account for nearly all of the hyperphagia, but not reduced energy expenditure, that leads to increased body mass seen in the global MC4R knock out mice[9, 10]. In addition, MC4R KO animals have altered meal structure and motivation to work for food reward in operant behavior paradigms, suggesting a coordinating role in food-seeking behavior in addition to body weight[11]. It is known that MC4R is expressed throughout the mouse cortex[12], including in the mPFC. Because of the mPFC role in body weight and food intake, we hypothesized that there is a subpopulation of MC4R expressing neurons in this region that coordinates the behavior functions of food-seeking or intake, which could provide new insight to the mechanisms underlying disordered eating behavior in health and disease.

## 2. Methods

### 2.1 Animals

Animals were housed in either the McLean Hospital Animal Care Facility, or the Beth Israel Deaconess Medical Center Animal Research Facility, with a 12h light/dark cycle, and given food and water *ad libitum*. All animals were individually housed for 1 week prior to feeding studies or other behavior testing. Animals were habituated to human handling during this time and body weight was measured daily throughout the duration of the behavioral studies. For animals injected with AAV-cre virus, the extent of the deletion was evaluated post-mortem, and only animals whose viral injection / MC4R deletion were confined to bilateral infralimbic cortex (IL) were used in the behavioral analysis, leading to exclusion of approximately ½ of the cohort (Fig. 2b).

### 2.2 Viruses

Adult male MC4R^lox/lox^ mice[10] were injected with were injected with pAAV.CMV.HI.eGFP-Cre.WPRE.SV40 (Addgene Cat. No. 105545-AAV9) into IL (bregma +0.175 AP, +/-0.03 ML, -0.03 DV) or injected with pAAV.hSyn.eGFP.WPRE.bGH (Addgene Cat. No. 105539-AAV9) into IL as controls. MC4R-2a-cre mice (JAX stock #030759)[13] were crossed to L10-GFP reporter mice[14] for use in imaging and electrophysiology experiments, and they were injected with AAV8-DIO-synaptophysin-YFP (Virovek, inc) for tracing experiments. *AgRP-IRES-Cre* mice[15] (Jackson Labs Stock 012899) were injected with AAV1-Syn-FLEX-GCaMP6s (Penn Vector Core, Addgene 100845) for photometry experiments.

### 2.3 Stereotaxic surgery and viral injections

For viral injections, mice were anaesthetized with ketamine/xylazine (100 and 10□mg/kg, respectively, i.p.) and then placed in a stereotaxic apparatus (David Kopf model 940). A pulled glass micropipette (20-40 μm diameter tip) was used for stereotaxic injections of AAV. Virus was injected into the IL-mPFC (25 nl/side; bregma +0.175 AP, +/-0.03 ML, -3.0 DV) or Arcuate (50 nl/side; bregma -1.5 AP, +/-0.25 ML, -5.8) by an air pressure system using picoliter air puffs through a solenoid valve (Clippard EV 24VDC) pulsed by a Grass S48 stimulator to control injection speed (40 nL/min). The pipette was removed 3 min post-injection followed by wound closure using tissue adhesive (3M Vetbond). An optic fiber (200 μm diameter, NA=0.39, metal ferrule, Thorlabs) was implanted in the IL (bregma +0.175 AP, +/-0.03 ML, -3.0 DV) and secured to the skull with dental cement. Subcutaneous injection of sustained release Meloxicam (4 mg/kg) was provided as postoperative care. The mouse was kept in a warm environment and closely monitored until resuming normal activity.

### 2.4 Behavior studies

#### 2.4.1 Home cage feeding

Animal food intake was analyzed by measuring the amount of food consumed every 12 hours of single housed animals (n=5 KO, n=6 control) for 2 weeks. Food was measured around the time of the light cycle change (ZT0 and 12). Time to approach and eat the food was manually scored.

#### 2.4.2 Open field

Animals were fasted for 16h overnight and then allowed to explore a dimly lit 40cm x 40cm arena for 10 minutes at ZT3-4. Motion (distance, velocity, and location) was tracked using Ethovision XT (Noldus).

#### 2.4.3 Novelty-suppressed feeding

Animals were fasted for 16h overnight and then introduced to a dimly lit 40cm x 40cm arena with a pre-weighed pellet of food in the center for a 10min epoch at ZT3. Motion (distance, velocity, and location) was tracked using Ethovision XT (Noldus), time to approach and consume the food was scored, and the food pellet was weighed.

### 2.5 Electrophysiology

Coronal brain slices from adult MC4Ra-Cre X L10GFP mice (male) containing medial prefrontal cortex (mPFC, 300 μm in thickness) were cut with a vibratome in cold oxygenated solution containing (in mM) 252 sucrose, 2.5 KCl, 5.0 MgSO_3_, 1.0 mM CaCl_2_, 26 NaHCO_3_, 1.25 NaH_2_PO_4_ and 10 glucose. Slices were kept in oxygenated artificial cerebrospinal fluid (ACSF) containing (in mM) 125 NaCl, 2.5 KCl, 2.5 CaCl_2_, 1.0 MgSO_4_, 26 NaHCO_3_, 1.25 NaH_2_PO_4_ and 10 glucose for at least one hour at room temperature before recording. MC4R-GFP positive neurons in the deep layer of infralimbic mPFC were visualized using differential interference contrast (DIC) optics (Zeiss 2FS microscope) and confirmed to express MC4R-GFP by observing fluorescence in response to blue LED illumination (excitation wavelength 470 nm, ThorLab). Membrane properties of MC4R-GFP neurons were assayed in whole-cell current-clamp recording mode in control ACSF containing NBQX (10 μM, AMPA receptor antagonist), D-AP5 (50 μM, NMDA receptor antagonist) and bicuculline (10 μM, GABA_A_ receptor antagonist) at temperature 32-34°C. Potassium-based intrapipette solution contained (in mM): 130 K-Gluconate, 5 KCl, 2.5 NaCl, 1.0 MgCl_2_, 0.2 EGTA, 10 HEPES, 2.0 ATP and 0.2 GTP. To examine the effect of THIQ (MC4R agonist, Tocris) and AgRP (MC4R inverse agonist, courtesy of Glenn Millhauser) on membrane properties of MC4R-GFP neurons, we continuously measured membrane potential and the number of action potentials generated by depolarizing current injections. Changes in the resting membrane potential and number of action potentials were assessed by comparing measurements during the pre-agonist baseline recording and those obtained 10 min after THIQ application.

### 2.6 Histology and Microscopy

Neurobiotin (6 mM) was added in the intrapipette solution to locate the recorded MC4R-GFP positive and negative neurons in the mPFC. After recordings, the slices containing recorded cells were fixed in the PBS containing 4% Paraformaldehyde overnight. Slices were washed in PBS and incubated in 0.1% Triton PBS containing streptavidin Alexa 568 conjugate (20 μg/ml, Molecular Probes) for two hours at room temperature, and washed again in PBS for three times (20 minutes each time). Finally, slices were mounted on gelatinized slides, dehydrated and coversliped. Vectashield antifading mounting medium (Vector Laboratory) was applied to slices to prevent fluorescence fading. Leica TCS SP8 confocal microscope (Leica) or Zeiss Axioskop 2 fluorescent microscope (Carl Zeiss) were used to capture images. Imaging data were processed and analyzed with ImageJ software (NIH).

### 2.7 Immunohistochemistry

Subjects were terminally anesthetized with 70% Ketamine and 10% xylazine and transcardially perfused with ice cold phosphate buffered saline (PBS) followed by ice cold 4% paraformaldehyde (PFA) in PBS. Following overnight fixation in 4% PFA at 4C, brains were transferred to a 30% sucrose solution at 4C until no longer floating. Brains were frozen in OCT and sectioned at 35 microns using a cryostat. Free-floating sections were washed 3 times in 0.1M PBS, then once in PBS with 0.05% Tween 20 (PBS-T). Sections were blocked for 1h at room temperature with 2% normal donkey serum in PBS-T, then incubated overnight at 4C in block with goat anti-AgRP, 1:1000 (catalog no. GT15023, Neuromics, Minneapolis, MN) and rabbit anti-POMC precursor 1:1000 (catalog no. H-029-30, Phoenix Pharmaceuticals). The following morning, sections were washed three times in PBS-T and incubated in block with AlexaFluor antibody 1:1000 (Molecular Probes, A-11055, A-31573) at room temperature for 2h in the dark. Sections were washed three times in PBS-T, then mounted on gelatin-coated slides, coverslipped with mounting media containing DAPI, and sealed with nail polish. Images were captured on Revolve ECHO microscope or on Olympus VS120 slide scanner microscope.

### 2.8 *In situ* hybridization

To examine gene expression for MC4R, tissue samples underwent single molecule fluorescent in situ hybridization. Anesthetized mice were decapitated, brains harvested, and flash frozen embedded in OCT on dry ice. Brains were stored at -80°C until cryostat sectioning, at which time they were equilibrated to -15°C in the Leica cryostat. Brains were coronally sectioned at 14 um and mounted onto Superfrost plus slides (Fisher), then air-dried at -15°C for 1h prior to storage in -80°C. MC4R probe and RNAscope fluorescent *in situ* hybridization multiplex kit was used according to the manufacturer instructions (Advanced Cell Diagnostics). Sections were imaged using an Olympus IX81 inverted fluorescence automated live cell microscope and processed with ImageJ software.

To examine gene expression for glutamate decarboxylases 1 and 2 (GAD1/2) and vesicular glutamate transporter 1, 2, and 3 (VGLUT 1/2), Molecular Instruments’ (Los Angeles, CA) *in situ* hybridization chain reaction protocol was used. Brains were stored and prepared similarly to those analyzed with RNAscope reagents. Brains were stored at -80□ until cryostat sectioning, at which time they were equilibrated to -15□ in the Epredia CryoStar NX70 Cryostat. Brains were coronally sectioned at 10 μm and mounted onto cold Superfrost Plus slides (Fisher). Slides were immediately kept on ice and later stored in -80□ freezer until further use. Reagents and probes were custom made by Molecular Instruments and used according to the manufacturer instructions. Sections were imaged using a Revolve fluorescent microscope or a Leica SP8 confocal. Images were processed with ImageJ and the plugin Bio-Format.

### 2.9 Statistical analysis

Data were analyzed using GraphPad Prism. Body weight and food intake were analyzed using two-way mixed measures ANOVAs and Bonferroni post hoc comparisons following significant interactions. All significance was defined as p<0.05.

## 3. Results

### 3.1 Identification of MC4R-expressing neurons in prefrontal cortex

Using a genetic cross of an MC4R-2a-cre mouse to an L10-GFP reporter mouse[13, 14], we found a dense population of MC4R-expressing neurons throughout the prefrontal cortex region (Fig. 1a). These are glutamatergic pyramidal neurons that co-express *SLC17A6*, the gene that encodes vesicular glutamate transporter 2 (vGlut2) and are found across the layers of the cortex (Fig. 1b). These neurons rarely express the other markers for glutamatergic vesicular packaging, vGlut1 or 3, and they do not express the markers for GABA (supplemental figure 1a). We investigated potential inputs from AgRP and POMC neurons of the arcuate nucleus of the hypothalamus to the prefrontal cortex, and found that they converged in the ventral part of the medial prefrontal cortex, the infralimbic region (IL) (Fig 1c). Therefore, we focused on MC4R-expressing neurons in the IL (IL^MC4R^) based on their proximity to peptidergic projections, and the region’s demonstrated involvement in goal-directed and habit-based activity, which may be relevant to hunger and feeding[16]. To identify the downstream projections of IL^MC4R^ neurons, we injected a virus containing a cre-dependent fluorescent marker, mCherry, into the IL of the MC4R-2a-cre mouse. IL^MC4R^ neurons projected most robustly to the medial septum, shell of the nucleus accumbens, lateral hypothalamus, and supramammillary nucleus, as well as a number of other regions (Fig. 1d, supplemental figure 1). Together, this combination of inputs from hypothalamic metabolic neurons to the IL region, and the robust projections from IL^MC4R^ neurons to regions known to be involved in motivated behavior, indicates that the IL^MC4R^ neurons may be an important node in the circuitry of food-seeking and/or intake behavior.

**Figure 1:**
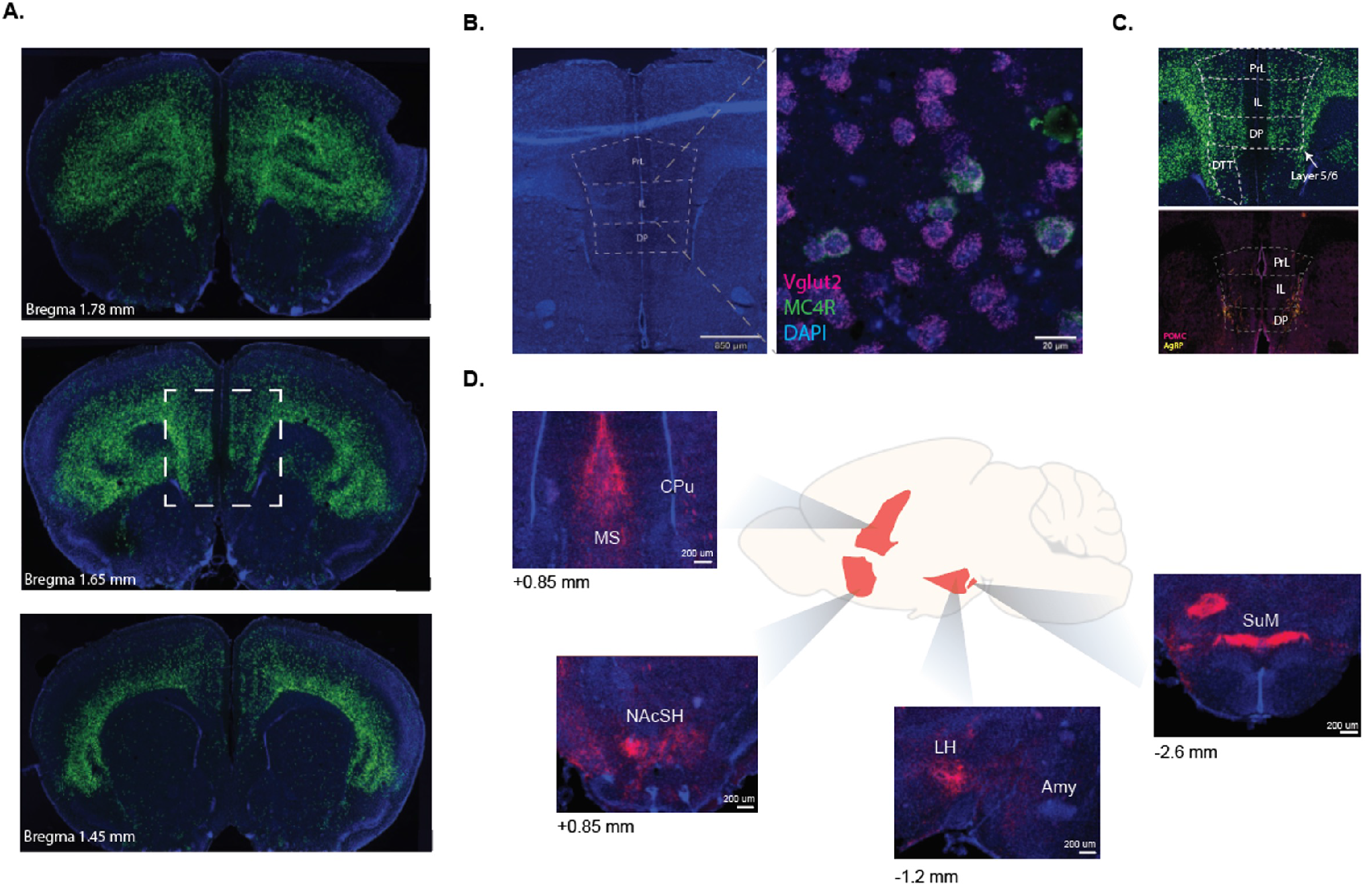
MC4R is expressed in a subpopulation of infra-limbic mPFC projection neurons that are glutamatergic and project downstream to regions implicated in food-motivated behavior. A. Melanocortin-4-receptor (MC4R) is expressed throughout the prefrontal cortex from bregma +1.78 to +1.45 mm in mice genetic cross of MC4R-2a-cre and fluorescent cre-dependent GFP reporter. Dotted line inset indicates subpopulation of MC4R-expressing neurons in the IL (IL^MC4R^); relevant for later images. B. *In situ* hybridization indicates colocalization of MC4R expressing neurons (green) with vesicular glutamate transporter 2 (magenta) in MC4R-2a-cre slice (bregma +1.55 mm). Image was taken at 10x and inset indicates a close-up at 40x. Dashed line shows pre-infralimbic (PrL), infra-limbic (IL), and DP (dorsal pedunculus) regions. C. Fluorescent image referring to the dotted inset in Fig.1A shows IL MC4R expression in MC4R-cre x L10 mice (bregma +1.67). Immunohistochemistry staining of AgRP and POMC projections performed in the same region on coronal slices from wild type mice. PrL (pre-limbic), IL (infralimbic), DP (dorsal pedunculus), DTT (dorsal tenia tecta). D. Schematic of downstream projections from mPFC^MC4R^ neurons, where colored regions indicate areas of greatest projection density. Insets show coronal slice immunohistochemistry. Presence of red indicates accumulation of fluorescent mCherry in projections from AAV-injections from mPFC^MC4R^ neurons. Medial septum (MS), caudate/putamen (CPu), nucleus accumbens shell (NacSh), lateral hypothalamus (LH), amygdala (Amy) supramammillary nucleus (SUM).

**Figure 2:**
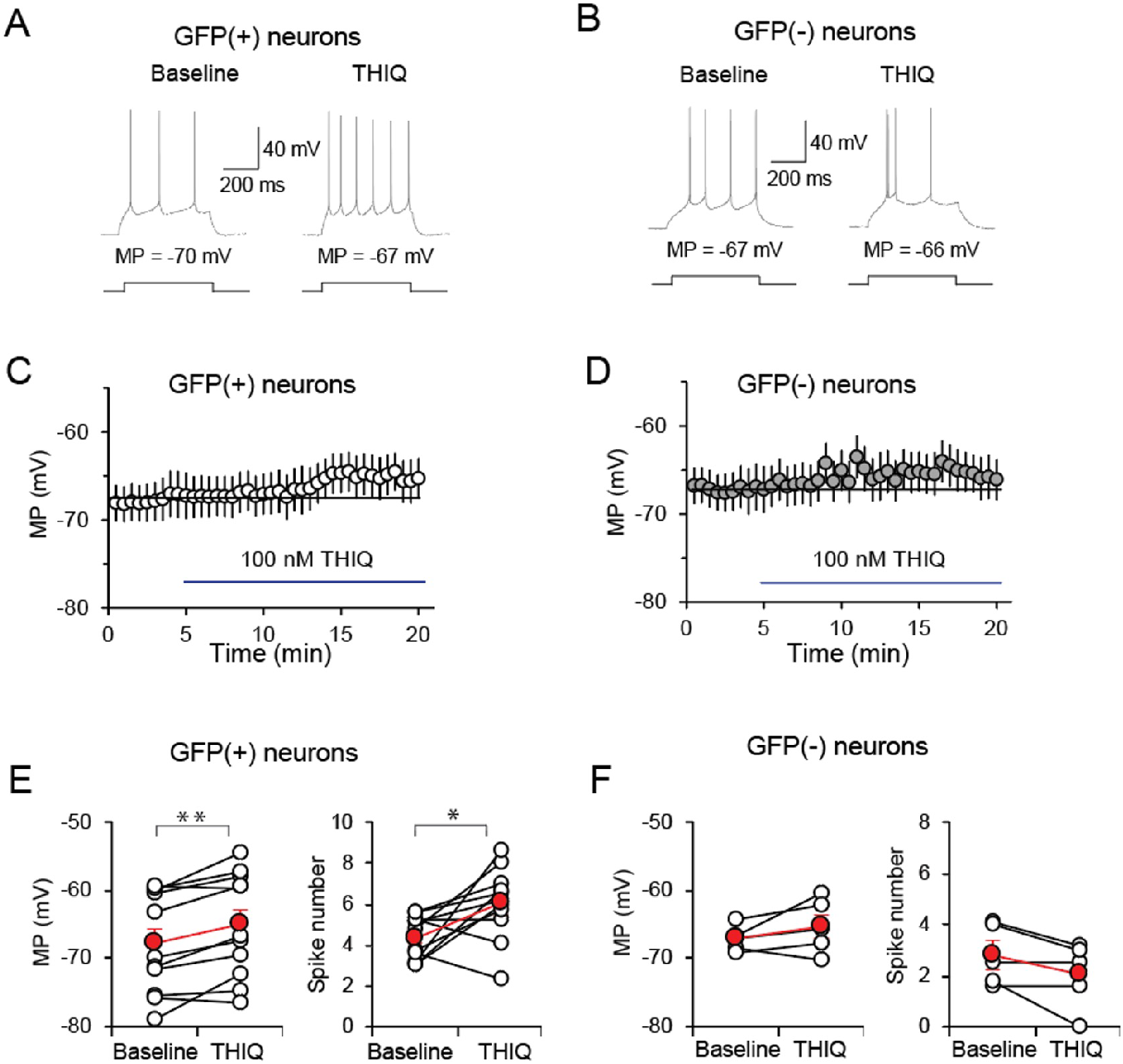
Depolarization of MC4R+ neurons in IL are mediated by MC4R. A. Example traces of action potentials recorded in MC4R-GFP positive (+) neurons were generated by depolarizing current injections (500 ms-long, once every 30 s) during pre-agonist recording (left) and 10 min after THIQ application (right). B. Example traces of action potentials recorded in MC4R-GFP negative (-) neurons were generated by depolarizing current injections (500 ms-long, once every 30 s) during pre-agonist recording (left) and 10 min after THIQ application (right). C. Resting membrane potential (MP) was continuously recorded in MC4R-GFP(+) neurons pre-treatment (for 5 min) and during bath application of MC4R agonist THIQ (100 nM). Data points in the panels represent averaged values of membrane potential in 30 s bins. Dashed lines show the averaged membrane potential level during pre-agonist baseline recording. D. Membrane potential (MP) was continuously recorded in MC4R-GFP (-) neurons for 5 min before and during bath application of MC4R agonist THIQ (100 nM). Data points in the panels represent averaged values of membrane potential in 30 s bins. Dashed lines show the averaged membrane potential level during pre-agonist baseline recording. E. Summary plot of the same experiments in panels A and C (GFP positive cells), showing averaged MP values (left) and the average number of action potentials (right) of 10 measurements in 5 minutes during baseline recording and 10 minutes after bath application of THIQ (paired t test; P = 0.003 and 0.025 for the measurement of MP and averaged number of AP, respectively, n = 11 GFP positive neurons from 7 mice). Open circles and red filled circles represent recordings of individual cell and averaged values, respectively. F. Summary plot of the same experiments in panels B and D (GFP negative cells) showing averaged MP values (left) and the average number of action potentials (right) of 10 measurements in 5 minutes during baseline recording and 10 minutes after bath application of THIQ (paired t test; P = 0.214 and 0.095 for the measurement of MP and averaged number of AP, respectively, n = 5 GFP negative neurons from 3 mice). Open circles and red filled circles represent recordings of individual cell and averaged values, respectively.

### 3.2 IL^MC4R^ are functionally responsive to melanocortinergic input

To determine whether the hypothalamic inputs to the IL region are functionally relevant, we tested whether IL neurons are responsive to the MC4R ligands produced by the hypothalamic feeding neurons. To do this, we recorded *in vitro* electrophysiological responses of MC4R-expressing (GFP+) and MC4R-non-expressing (GFP-) IL neurons to exogenously applied THIQ, an MC4R specific agonist, to cortical slices obtained from MC4R-2a-cre x L10-GFP reporter mice. (supplemental Fig. 2a). The average pre-agonist baseline was approximately the same for both GFP+ and GFP-neurons (Fig. 2c,d), but application of THIQ lead to a relative depolarization of resting membrane potential in GFP+ neurons (Fig. 2c,e), that was not seen in GFP-neurons (Fig. 2d,f). In addition, there was a significant increase in response to THIQ in depolarization-induced action potential firing frequency in GFP+ neurons (Fig. 2a,e), not seen in GFP-neurons (Fig. 2b,f). Since THIQ mimics the stimulatory input of αMSH produced by POMC neurons on MC4R, these data suggest that satiety-inducing melanocortinergic neuropeptide signals act via MC4R to depolarize and increase spiking activity in mPFC^MC4R^ neurons, suggesting that stimulation of the IL^MC4R^ by αMSH may be important for food-seeking or intake behavior.

### 3.3 IL^MC4R^ are important to maintain body weight and food intake

Dysregulated food-seeking behavior can cause a change in food intake. To identify whether MC4R expression in the IL region contributed to food intake and downstream metabolic changes, we created a conditional viral-mediated and cre-dependent deletion of MC4R in IL (IL^MC4R^ KO) (Fig 3a,b). We compared home cage spontaneous food intake between the IL^MC4R^ KO and control groups injected with AAV-GFP. We measured food intake twice daily for two weeks after viral injections, during which time the body weight between the two groups was not significantly different. Among the IL^MC4R^ KO mice, only those for whom injections (and deletion) were restricted to the IL region showed a significant increase in food intake (approximately 0.5g more standard chow per night) compared to control mice (Fig 3c), indicating that deletion of MC4R from the IL only has an effect on food intake. Interestingly, animals with MC4R deleted from a larger area of the mPFC including both the prelimbic and infralimbic regions showed no change in food intake behavior (supplemental Fig 3a). In all KO animals with bilateral deletion covering the IL, starting approximately 3 weeks post MC4R viral mediated deletion, we observed a robust and significant increase in body weight (Fig 3d). Intriguingly, animals with KO that also hit the more dorsal prelimbic region demonstrated a trend to decreased bodyweight (supplemental Fig 3b). This suggests that the regions of the mPFC have different roles in energy regulation, and that the hyperphagia induced by IL^MC4R^ deletion is not compensated for by changes in energy expenditure that would limit the development of obesity.

**Figure 3:**
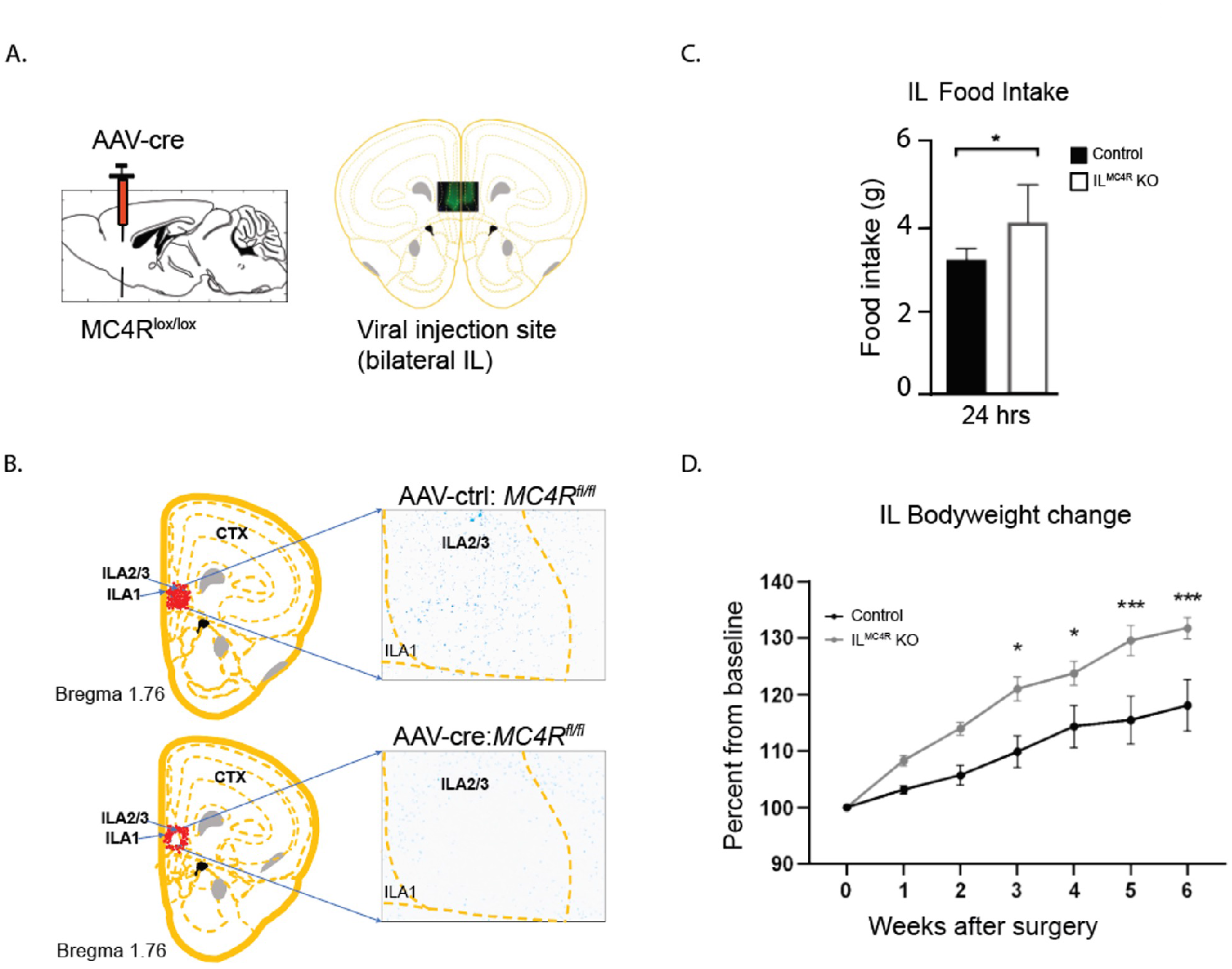
Viral cre-mediated KO of MC4R in IL^MC4R^ neurons leads to an increase in food intake and body weight. A. (Left) Schematic of MC4R deletion in IL^MC4R^ neurons by injection of adeno-associated virus into IL of MC4R^lox/lox^ mice. (Right) Fluorescent image of bilateral viral injection into the IL overlaid on a coronal tracing of the same region (bregma +1.67 mm). B. Diagram of injection site and the average MC4R expression pattern around IL before and after injection. Insets show conditional knock-out of MC4R mRNA (blue) in IL using RNAscope. Image is inverted for ease of visualization. C. MC4R^lox/lox^ mice injected with AAV-cre (n=5) restricted to the IL ate approximately 0.5g (∼12.5%) more on average over the course of 2 weeks prior to onset of body weight gain than those injected with AAV-GFP (n=6), (2-tailed unpaired t-test *p<0.05). D. MC4R^lox/lox^ mice injected with AAV-cre (n=8) restricted to the IL gained more weight than those injected with AAV-GFP (n=7) following surgery (two-way mixed measures ANOVA, treatment by week interaction, F(6,78)=4.736, p=0.0004. Bonferroni post hoc comparison, *p<0.05 ***p<0.001 between groups).

### 3.4 IL^MC4R^ are relevant to food-seeking, but not general exploratory behavior

The mPFC in rodents is critical for coordinating goal-directed and habitual behavior[2]. In studies of reward-seeking, the subregion of the IL has been shown to be important for inhibition or disinhibition of action[17, 18], which might be relevant to both food-seeking and food consumption. To consider the contribution of IL^MC4R^ to this type of executive function behavior, we examined food-seeking in a novel environment. Importantly, the IL^MC4R^ KO animals showed no difference in exploratory behavior in the novel environment compared to intact animals (Fig 4a), so there was no effect of the IL^MC4R^ KO on locomotor activity or anxiety-associated inhibition of exploration. In addition, the groups showed no difference in food-approach or consummatory behavior in the home cage in a 20-minute bout after overnight food deprivation (Fig 4b), indicating food-seeking and intake in the acute setting are intact. However, when presented with a piece of food in the center of the novel environment after an overnight fast, the IL^MC4R^ KO animals took double the amount of time to approach the food as well as to begin consuming it (Fig 4c). There was no significant difference in the amount consumed in a 20 minute time period between groups, though both groups of animals did eat more in that time period in the home cage compared to the novel setting (Fig 4c). These findings implicate the involvement of executive function, as the animal must choose to interact with food in the novel environment, and the IL^MC4R^ appears to be important for this behavior function.

**Figure 4:**
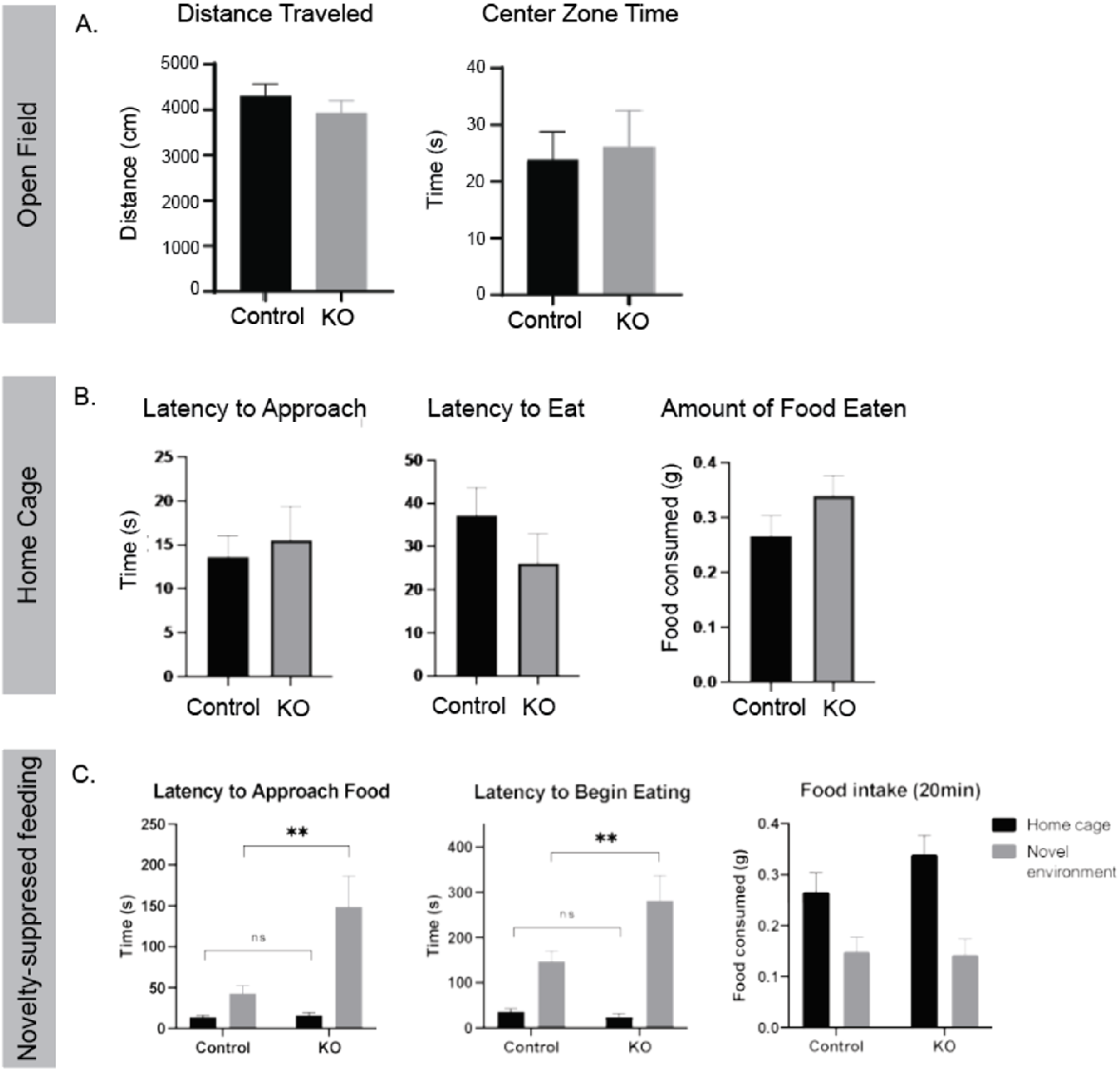
Deletion of IL^MC4R^ leads to increased latency in approaching food in novel environments but no change in food interactions in home cage. A. No group differences were observed in distance traveled (left) or total time spent in the center zone of open field (right) for IL^MC4R^ KO mice and control mice (Student’s t-tests, p>0.05). Error bars ± S.E.M. (KO n =10, ctrl n=9). B. AAV-cre-injected IL^MC4R^ (KO) and AAV-ctrl (ctrl) littermates were fasted overnight and then given access to food in their home cage. IL^MC4R^ KO mice (n=10) show no significant difference to approach (left) and eat (center) food in home cage than control mice (n=9) (two-way mixed measures ANOVAs, treatment by environment interactions, F(1,17)=6.261, p=0.0228 and F(1,15)=6.325, p=0.0238. Bonferroni post hoc comparisons, **p<0.01). Error bars ± S.E.M. (Right) IL^MC4R^ KO mice (n=10) and control mice (n=9) show no significant difference in the amount of food consumed when in their home cage. C. AAV-cre-injected IL^MC4R^ (KO) and AAV-ctrl (ctrl) littermates were fasted overnight and then given access to food, either in their home cage or in the center of a novel environment. IL^MC4R^ KO mice (n=10) take longer to approach (left) and eat (center) food in a novel environment than control mice (n=9), but show no difference in home cage feeding (two-way mixed measures ANOVAs, treatment by environment interactions, F(1,17)=6.261, p=0.0228 and F(1,15)=6.325, p=0.0238. Bonferroni post hoc comparisons, **p<0.01). Error bars ± S.E.M. (Right) IL^MC4R^ KO mice (n=10) and control mice (n=9) ate significantly more in the home cage than the novel environment but showed no differences in food intake within the IL^MC4R^ KO and control groups.

## 4. Discussion

In this study, we show that there is a novel population of glutamatergic neurons in the IL that express MC4R, and deletion of MC4R from only these neurons significantly alters food intake and body weight. We demonstrate that interruption of the IL^MC4R^ signal in male mice acutely increases food intake behavior and leads to longer term increases in body weight. The weight gain of the IL^MC4R^ KO is markedly greater than the minimal weight gain seen in other region-specific conditional knockouts of MC4R outside of the paraventricular hypothalamus (PVH), such as the medial amygdala and lateral amygdala, regions of the brain that are involved in body weight, but where deletion of the MC4R has no effect on either food intake or body weight. Deletion of the MC4R from the nucleus of the solitary tract was shown to lead to ∼10% reduction in body weight, with no effect on food intake[10]. Other prior work has shown that manipulation (re-expression or deletion) of PVH MC4R alone accounts for 60% of the obesity and 100% of the hyperphagia[9, 10] and is 50% of that of whole body MC4R knockouts[19]. Taken together, this suggests a primary etiologic role for the IL in the regulation of food intake. In addition, the 10-20% increase in body weight seen in the IL^MC4R^ KO accounts for the difference in body weight and food intake between vGlut2-mediated MC4R deletion and Sim1-mediated PVH MC4R deletion[10], indicating this novel glutamatergic population fills that gap, and is important for connecting hyperphagia directly to resultant body weight change. While not examined here, it will be important to separately investigate the role of IL^MC4R^ in energy expenditure.

Importantly, we show that an increase in food intake is seen only when deletion of MC4R is restricted to the ventral region of the mPFC, not when the deletion includes both prelimbic and infralimbic regions. This reinforces previous research showing that manipulation of neurons in the whole mPFC enhances food seeking behavior, but does not change food intake[20, 21]. Furthermore, these results suggest that food-intake related behavior, potentially related to cognitive functions, are differentially regulated by mPFC subregions, conceptually consistent with results supporting distinct roles for mPFC subregions important for the expression of appetitive vs. aversive instrumental behaviors[16–18, 22, 23]. Much work remains to be done to understand the distinct role of IL in appetitive responses, whether conditioned or innate. We hypothesize that these changes reflect an important role for IL^MC4R^ in feeding-related executive function.

There are three core executive functions coordinated in the mPFC as defined in humans: inhibition, working memory, and cognitive flexibility[24]. In rodents these functions are also attributed to the medial prefrontal cortex, but the anatomic distinctions of the region carry different functionality. The IL region is important in habitual processes, while the more dorsal prelimbic region (PL) is necessary for goal-directed behavior[25]. In addition to habit, the IL region has been shown to be involved some specific elements of executive function, including reward seeking as well as appetitive behavior inhibition[17, 26, 27]. Inhibition can be described as self-control of action-oriented behavior, and also interference control, with selective attention biased toward a response to one out of multiple sensory inputs.

In our studies, the executive function deficits we observe could involve both. Behavior inhibition is seen through an impairment in the discontinuation of food intake (habit-control), which leads to hyperphagia and increased body weight in the home cage. Alternatively, selective attention attributed to one of two potentially conflicting salient stimuli (center of open field and access to food) implies a change in interference control. We show that the animals with MC4R deletion from IL interact acutely in the home environment with food in a manner similar to animals with intact IL^MC4R^ in the setting of a fast: both groups of animals very quickly approach and consume the food presented, indicating that the valuation of the food in the hunger state, and home cage food-seeking behavior are intact. Additionally, both groups behave the same in the open field environment, indicating that the response to the novel space, often considered a readout of anxiety-like behavior, is not different between groups. However, when we provide fasted animals with these inputs together – a food pellet in the center of an open field - we find that the groups diverge. The IL^MC4R^ KO animals take longer to approach and to consume the food, indicating the balance of these two inputs that leads to a particular expected behavior output is impaired, and the input of potential threat of the center of a novel space is favored, for a period of time, over the hunger-related input. Future studies will be needed to investigate if the IL^MC4R^ impairment can be dissociated between food-seeking behavior (goal-directed, selective attention) or to discontinuation of food intake (habit-based override).

Finally, we show that IL^MC4R^ neurons are uniquely responsive to melanocortinergic stimulation, and are distinguished by being localized in a section of the medial prefrontal cortex that receives inputs from the hypothalamic melanocortin peptide producing neurons. This could be part of a metabolically sensitive pathway whereby information about body nutrient status influences body and behavior function, as is known to occur in the hypothalamus[28]. Another putative explanation is that IL^MC4R^ neurons are part of a context-dependent pathway, whereby external information is encoded and is integrated further downstream, which is a known function of the IL[16]. A final alternative is that MC4R expression could provide the molecular definition for a neural ensemble that is specifically responsive to natural food reward, as it has been shown in prior work that different IL neurons are activated by different forms of reward[23, 29]. Future work will be required to investigate if the peptidergic input is synaptically derived, and part of a functional direct circuit from the hypothalamic melanocortinergic neurons.

To understand the specificity of this MC4R population for food-related decision making, it will be important to disentangle food motivation from other forms of natural reward motivation. Because so many learning and motivation paradigms rely on food restriction for training, and because AgRP stimulation has been shown to drive non-food related compulsive seeking behavior[30, 31], it remains unclear if an MC4R-related signal is solely a metabolism-related stimulus. The mPFC has been implicated in the consummation of water and sexual behavior[5, 32], so it will also be important to identify the contributions of the IL^MC4R^ neurons in driving these consummatory goal-oriented behaviors. Also important for future study is to investigate these neurons in females, given the sex differences in symptoms and prevalence of various eating disorders and pathology related to hyperphagia in humans. Energy state-related signals and energy metabolism are different between males and females, from increased baseline leptin signal found in females to decreased food consumed in the setting of a fast[33]. Given the importance of sex as a biological variable, in both metabolism and motivated learning[34, 35], the role of sex hormones in metabolism [36], the association of MC4R mutations with more profound obesity in females[37] and more recently with stress-related metabolic outcomes[38], it will be relevant to investigate this circuit and related behavior in female mice, and take hormone cycle dynamics into account.

## Conclusions

Our findings show that the IL^MC4R^ neurons are functionally responsive to melanocortinergic peptides, and that MC4R expression in the IL is necessary for normal food-seeking and food intake that are required to maintain appropriate body weight. These data support the idea that melanocortinergic input may significantly contribute to executive function, and that integration of metabolism and executive function may be an important neural substrate mediating both systemic metabolic outcomes and behavior related to food intake, or other sensory information integration, in psychiatric disorders.

## Abbreviations

αMSH: alpha melanocyte stimulating hormone
AgRP: agouti related peptite
IL: infralimbic cortex
KO: knockout
mPFC: medial prefrontal cortex
MC4R: melanocortin 4 receptor
PL: prelimbic cortex
POMC: proopiomelanocortin

## Figure Captions

**Supplemental Table 1:**
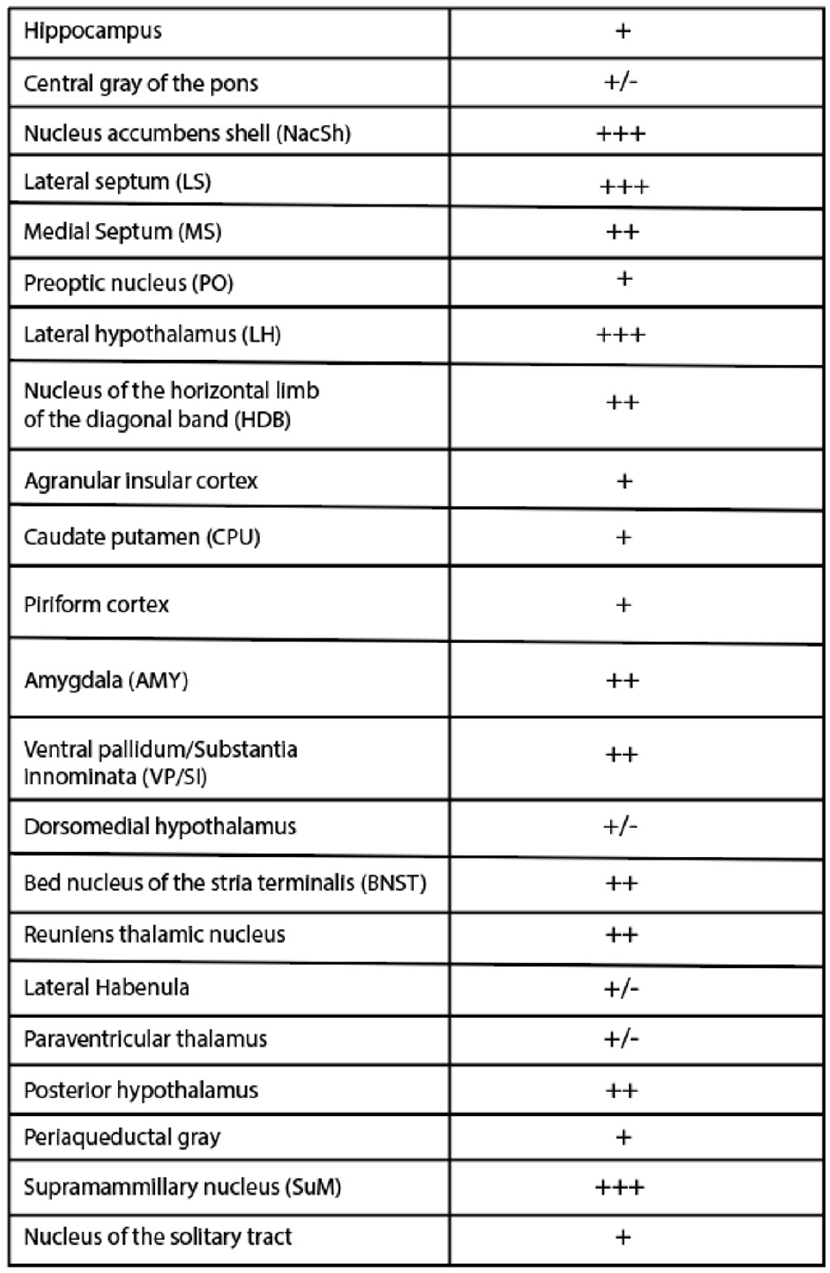
Projection density of mPFC^MC4R^ neurons found throughout the brain. A. Table of regions throughout the brain where projections of mPFC^MC4R^ neurons are found. Projection density is categorized as low (+), intermediate (++), and high (+++).

**Supplemental Figure 1:**
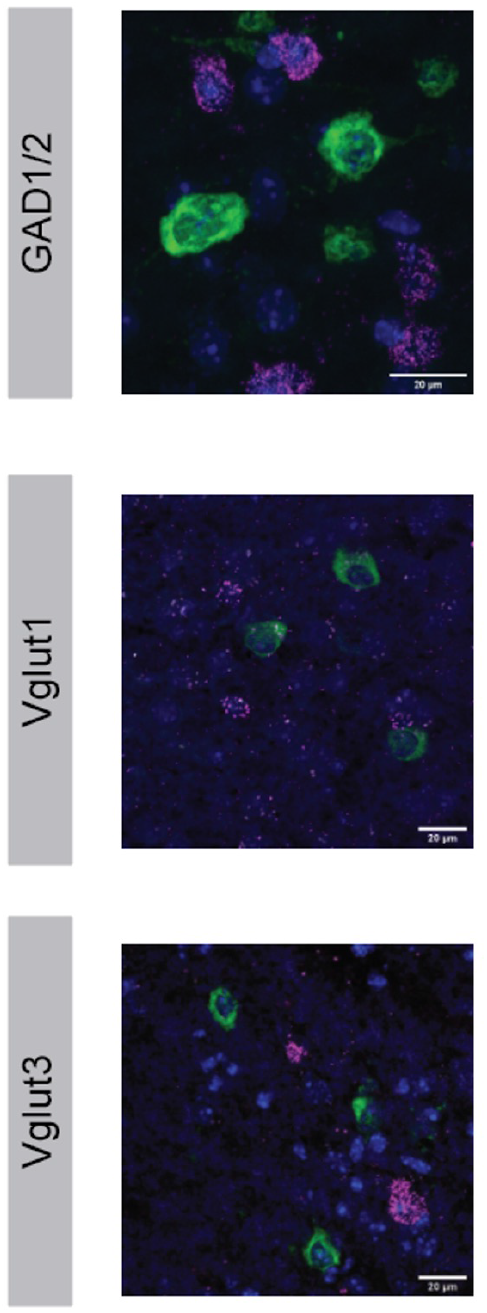
IL^MC4R^ neurons are not gaba-ergic and rarely express markers for vGlut1 and vGlut3. A. In-situ hybridization indicates no colocalization of MC4R expressing neurons with (top) gabaergic markers glutamic acid decarboxylase 1 and 2, (center) vesicular glutamate transporter 1, (bottom) vesicular glutamate transporter 3. Images were taken on confocal at 40x and processed in ImageJ.

**Supplemental Figure 2:**
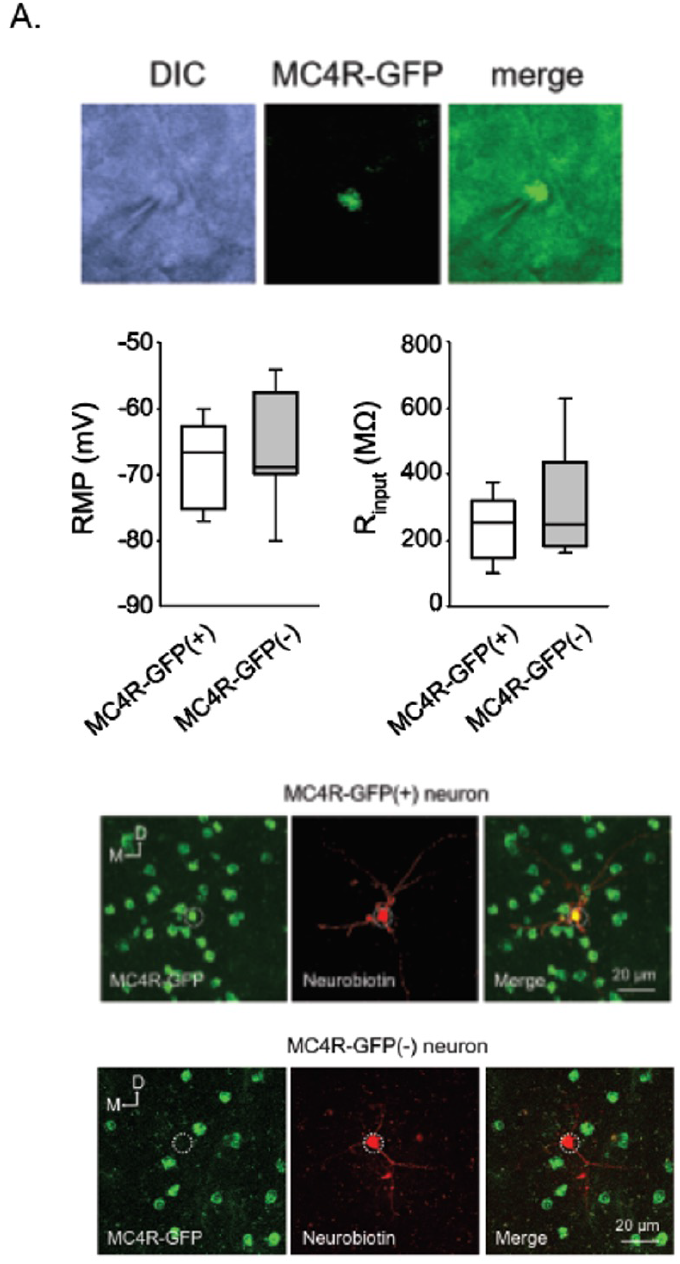
Validation of MC4R-GFP (-) neurons in the IL in normally fed male mice. A. Top, microscopic images of a recorded MC4R-GFP positive neuron in the IL (DIC image, left; MC4R-GFP fluorescence, middle; merged, right). Center right, box plot shows 25%-75% percentile, median, minimal and maximal value of the resting membrane potential (RMP) of MC4R-GFP(+) and MC4R-GFP(-) neurons. No difference between the two groups, unpaired t test, P = 0.51 (n = 28 MC4R-GFP positive neurons from 11 mice versus 11 MC4R-GFP negative neurons from 3 mice); center left, box plot shows 25%-75% percentile, median, minimal and maximal value of input resistance (Rinput) of the recorded neurons as in left. Unpaired t test, P = 0.19 between the two groups. Bottom, confocal stack images show an example of recorded MC4R-GFP(+) neurons and MC4R-GFP(-) neurons identified by colocalization (right, merged) of MC4R-GFP fluorescence (left, green) with intracellularly delivered Neurobiotin (red, middle) in the recorded neurons. The white circles indicate the soma location of recorded neurons.

**Supplemental Figure 3:**
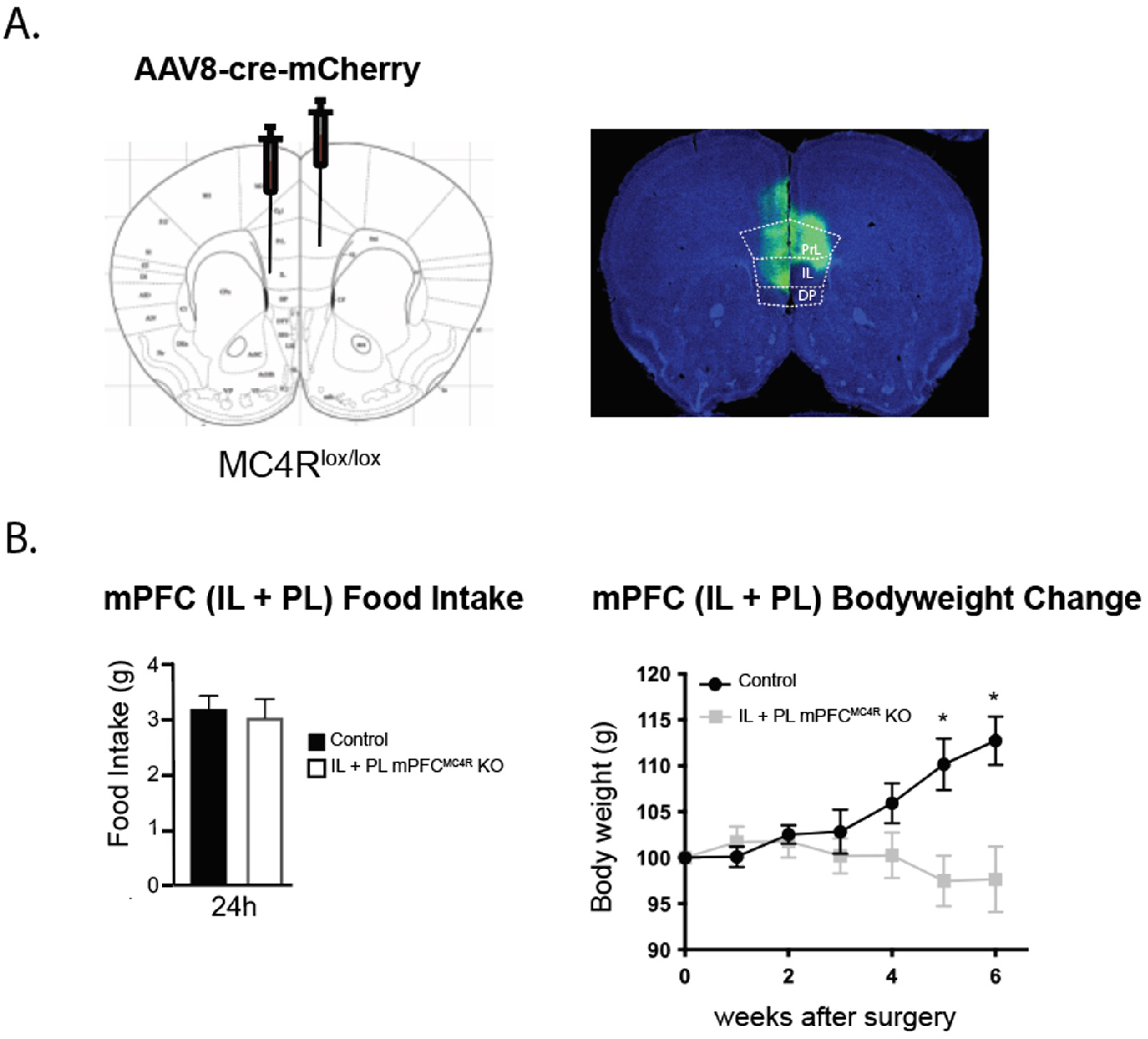
Increase in food intake and body weight is observed when conditional KO of MC4R is constrained to the IL region but not whole mPFC. A. Schematic of AAV cre-mediated KO of MC4R in the whole mPFC (IL and PL) of MC4R^lox/lox^ mice (left). Example of slice from mouse with PL + IL cre-injection (right). B. MC4R^lox/lox^ injected with AAV-cre (n=4) in PL+IL showed no difference in average food intake, and increased body weight, when compared to MC4R^lox/lox^ injected with AAV-GFP control virus (n=7).

## Acknowlegements

This work was supported by funding from NIH (UL1 TR001102, K08 DK118201) and the McLean Maria Lorenz Pope fellowship to RAR, as well as NIH awards (P50-MH115874, R01-MH108665), and the Frazier Institute at McLean Hospital to KJR.

The authors wish to thank Brad Lowell, Gary Schwartz, Jelena Radulovic, Saleem Nicola, Mark Baxter, Katherine Nautiyal, Alex Harris, and Natalie Rasgon for helpful input preparing the manuscript.

